# Suppression of alpha-band power underlies exogenous attention to emotional distractors

**DOI:** 10.1101/2021.02.22.432307

**Authors:** Lydia Arana, María Melcón, Dominique Kessel, Sandra Hoyos, Jacobo Albert, Luis Carretié, Almudena Capilla

**Affiliations:** Departamento de Psicología Biológica y de la Salud, Facultad de Psicología, Universidad Autónoma de Madrid, Spain; Departamento de Neurociencia y Aprendizaje, Universidad Católica de Uruguay, Uruguay

**Keywords:** Alpha oscillations, EEG, emotional distractor, exogenous attention, visuospatial

## Abstract

Alpha-band oscillations (8-14 Hz) are essential for attention and perception processes by facilitating the selection of relevant information. Directing visuospatial endogenous (voluntary) attention to a given location consistently results in a power suppression of alpha activity over occipito-parietal areas contralateral to the attended visual field. In contrast, the neural oscillatory dynamics underlying the involuntary capture of attention, or exogenous attention, are currently under debate. By exploiting the inherent capacity of emotionally salient visual stimuli to capture attention, we aimed to investigate whether exogenous attention is characterized by either a reduction or an increase in alpha-band activity. Electroencephalographic activity was recorded while participants completed a Posner visuospatial cueing task, in which a lateralized image with either positive, negative, or neutral emotional content competed with a target stimulus presented in the opposite hemifield. Compared with trials with no distractors, alpha power was reduced over occipital regions contralateral to distracting images. This reduction of alpha activity turned out to be functionally relevant, as it correlated with impaired behavioural performance on the ongoing task and was enhanced for distractors with negative valence. Taken together, our results demonstrate that visuospatial exogenous attention is characterized by a suppression of alpha-band activity contralateral to distractor location, similar to the oscillatory underpinnings of endogenous attention. Further, these results highlight the key role of exogenous attention as an adaptive mechanism for the efficient detection of biologically salient stimuli.

**Highlights:**

- Exogenous attention is indexed by alpha suppression contralateral to distractors.
- Alpha power decrease is enhanced by distractors with negative emotional valence.
- Lower levels of alpha power correlate with poorer task performance accuracy.
- The negativity bias in exogenous attention might reflect an adaptive mechanism.

## Introduction

The human brain is constantly exposed to a multitude of stimuli. Hence, in order to discriminate relevant from unnecessary information, an attention system becomes essential. Stimulus selection is accomplished according to internal goals or automatically triggered by certain stimuli, namely endogenous and exogenous attention, respectively (Chica et al., 2011; Corbetta et al., 2008). On the one hand, given that our processing resources are limited, endogenous attention is responsible for ignoring distractors in favour of the ongoing task (Lavie, 2005). On the other hand, if distractor stimuli reach a given saliency threshold, exogenous attention may be captured and cognitive resources diverted to them (Koster et al., 2004).

Alpha-band oscillations (8-14 Hz) are considered to be crucial for attention and perception, by facilitating the selection of relevant information and inhibiting irrelevant distractors (Foxe & Snyder, 2011; Jensen et al., 2014; Jensen & Mazaheri, 2010; Klimesch, 2012). Consistent evidence supports these theories, showing that modulations of alpha activity correlate with the direction of visuospatial attention. In particular, alpha-band power has been found to decrease over occipito-parietal areas contralateral to the attended visual field and/or to increase ipsilaterally (Capilla et al., 2014; Händel et al., 2011; Siegel et al., 2008; Thut et al., 2006; Worden et al., 2000; Wöstmann et al., 2019). In addition, a number of recent studies employing multivariate approaches have shown that selective topography of alpha oscillations precisely tracks the temporal dynamics of visuospatial orienting (Foster et al., 2017; Samaha et al., 2016). Importantly, the modulation of alpha power has an impact on behavioural performance, by enhancing the perception of the target (Capilla et al., 2014; Kelly et al., 2009; Thut et al., 2006) and disrupting the processing of distractors (Payne et al., 2013; Zumer et al., 2014). Furthermore, it has been convincingly demonstrated that alpha-band fluctuations do not only predict the locus of visuospatial attention but causally drive attention and visual perception (Bagherzadeh et al., 2020; Romei et al., 2010; Thut et al., 2011).

It is worth noting that majority of research has focused on endogenous attention, whilst the oscillatory neural dynamics of exogenous attention have been largely neglected. First evidences of modulations of alpha-band oscillations under exogenous attention requirements are quite recent and arise from cross-modal experiments, wherein peripheral non-predictive (i.e., 50% valid) auditory cues are used to involuntarily orient visuospatial attention (Feng et al., 2017; Keefe & Störmer, 2021; Störmer et al., 2016; Weise et al., 2020). Overall, these studies demonstrate that exogenous cues trigger an oscillatory mechanism within the alpha frequency band and enhance subsequent visual processing, resembling the positive influence of endogenous attention on perception.

However, although all of these studies agree on the relevance of the alpha rhythm, there are substantial discrepancies between them in terms of the pattern of power modulation found. In particular, two studies have shown that alpha power decreases over occipito-parietal regions, especially in the hemisphere contralateral to the cued side (Feng et al., 2017; Störmer et al., 2016), whereas another found that alpha-band power increases, rather than decreases, in the hemisphere ipsilateral to the auditory cue (Weise et al., 2020). A fourth study only reported lateralization effects (contra – ipsilateral difference), so the net effect of each hemisphere cannot be estimated (Keefe & Störmer, 2021). This discrepancy in the pattern of alpha power modulation is crucial for the interpretation of the mechanisms underlying visuospatial exogenous attention. Whilst a contralateral decrease in alpha-band power would imply the facilitation of stimulus processing at attended locations, an ipsilateral alpha increase would indicate the inhibition of the unattended visual field (Foxe & Snyder, 2011; Jensen et al., 2014; Jensen & Mazaheri, 2010; Klimesch, 2012). Weise and colleagues (2020) suggest that the discrepant results of their study could be due to the fact that they included a distraction task in addition to the spatial cueing task.

The present study sought to shed light on these conflicting findings, by using a type of stimulus that inherently induces distraction: emotional images. It has been pointed out that emotion-laden distractors grab exogenous attention to a greater extent than neutral stimuli (Carretié, 2014). This is not surprising from an evolutionary perspective, given the importance of detecting biologically salient stimuli such as danger, food, or mating partners, which are, by definition, emotional. We thus harnessed the engaging property of emotional images to investigate the oscillatory mechanisms underlying visuospatial exogenous attention.

We recorded electroencephalographic (EEG) activity while participants performed a Posner cueing task. A visual predictive cue indicated with a validity of 100% the location of an upcoming target, thus ensuring that endogenous attention was oriented to it. An emotional or non-emotional distractor image appeared simultaneously with the target in the uncued visual field. If exogenous attention were effectively captured by emotion-laden images, we would expect a suppression of alpha-band power over posterior regions contralateral to the distractor location. Alternatively, if task-irrelevant distractors were to be inhibited, we would expect an increase in contralateral alpha power. Our results support the former hypothesis, showing that visuospatial exogenous attention is characterized by a suppression of alpha power that is more prominent over contralateral occipital sites and for distractors with negative valence. This suggests that the desynchronization (reduction) of alpha-band activity indexes the degree of attentional capture and the resources diverted to distractor stimuli, with the consequent negative impact on behavioural performance.

## Methods

### Participants

Thirty-six volunteers took part in this study (19 females, 29 right-handed, 21.6 ± 3.5 years old, mean ± SD), although one was excluded from EEG analysis as explained below. They all reported normal or corrected-to-normal visual acuity. Before participating in the experiment, they provided informed written consent. The study was approved by the Ethics Committee of the *Universidad Autónoma de Madrid* and conducted in compliance with the declaration of Helsinki.

### Stimuli and procedure

Participants completed the experimental task while seated 1 m away from the screen inside an electrically shielded, sound-attenuated room. The task was programmed using Psychtoolbox (v3.0; Brainard, 1997) in Matlab (The MathWorks) and presented through a RGB projector on a back-projection screen.

The task was divided into four experimental blocks of 105 trials each. The structure of a trial is depicted in Figure 1. A central fixation cross was presented on a grey background, subtending 2.3 × 2.3° of visual angle. After a variable interval ranging from to 1 s, an arrow (0.6 × 0.6°) was appended to the fixation cross for 0.1 s, pointing to either the left or the right lower quadrant. This cue indicated with 100% probability the subsequent location of the target stimulus. Participants were instructed to pay covert attention (i.e., without moving their eyes) to the cued location.

**Figure 1.**
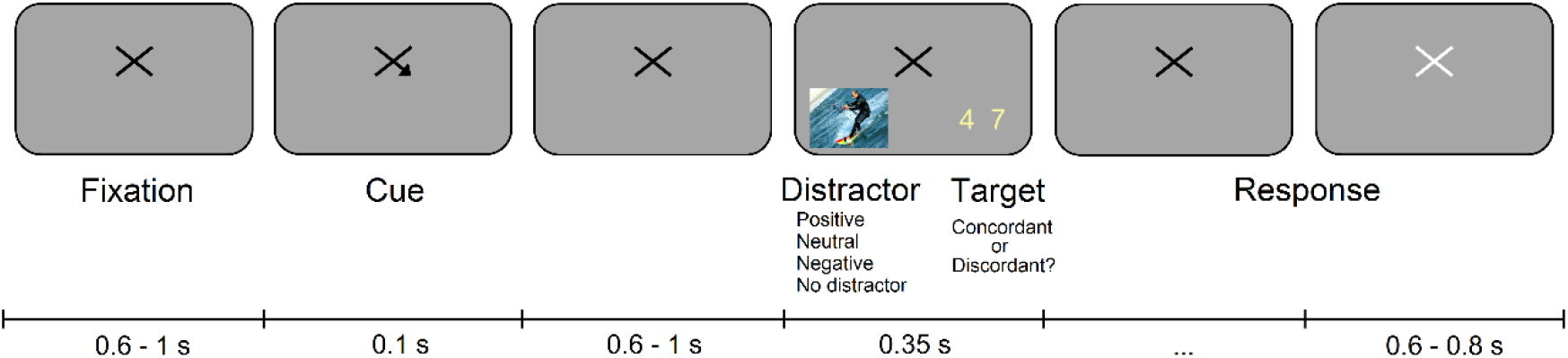
Experimental task. A spatial cue indicated with a probability of 100% the location of an upcoming target stimulus (i.e., a pair a numbers). Participants performed a digit-parity task while an emotional distractor appeared in the uncued hemifield. No distractors were presented in the control block.

After a variable delay of 0.6 to 1 s, a composite target stimulus composed of a pair of yellow numbers was displayed on the screen for 0.35 s. The size of the digits was 1.7 × 2.9° (width x height), and they were presented in either the left or the right lower quadrant at 10.8 × 9.4° from fixation. Participants performed a digit-parity task during which they were required to report if the pair of numbers was concordant (i.e., both digits odd or both even) or discordant (i.e., one odd and one even), by pressing a key as accurately and fast as possible. After providing the motor response, the fixation cross turned white for 0.6-0.8 s, indicating the beginning of the next trial. Participants were encouraged to use the time between trials to blink if necessary.

In three out of the four experimental blocks, a distractor stimulus appeared in the uncued location at the same time as the target stimulus. In the remaining block no distractor stimuli were presented, thus acting as a control. The order of presentation of the blocks was randomized across participants. Distracting images were obtained from the EmoMadrid emotional pictures database (Carretié et al., 2019), whose normative data range from −2 to +2 in both emotional arousal and valence dimensions. We selected three groups of 35 images with positive, neutral, and negative valence. Positive and negative images were matched in absolute valence (positive: |1.29| ±.148; negative: |-1.28| ± .157; t_68_ = .340, p = .735), whereas the valence of the neutral images did not significantly differ from zero (-.001 ± .130; t_34_ = -.069, p = .945). In a similar way, arousal was matched for emotion-laden images (positive: 1.27 ± .221; negative: 1.24 ± .190; t_68_ = .586, p = .559) and near zero for neutral images (-.001 ±.154; t_34_ = -.052, p = .959).

The size of the distracting images was 14.6 × 11.0° and they were positioned at a visual angle of 10.8 × 9.4°. On each trial, one image was pseudo-randomly chosen, with the condition that every image appeared three times throughout the whole experiment, whilst images with the same valence did not appear on more than two consecutive trials.

### Self-reported emotional valence and arousal

At the end of the recording session, participants completed a bidimensional scaling test for each of the 105 images, assessing both their valence and arousal. This last test aimed to confirm that *a-priori* categorization of stimuli as positive, negative, and neutral was valid in our sample. The effect of Distractor (positive, negative, neutral) on valence and arousal was statistically tested by means of a one-way repeated measures analysis of variance (ANOVA). We also conducted a series of t-tests to statistically check whether positive and negative images differed in absolute valence and arousal, and whether neutral images differed from zero in emotional valence and arousal. Significance threshold was adjusted by Bonferroni correction for multiple comparisons. All statistical analyses were performed using IBM SPSS Statistics 26.

### Behavioural analysis

Behavioural analyses were carried out to verify that distractor stimuli captured exogenous attention. If this were the case, we would expect lower accuracy and longer response times in the digit-parity task. Trials contaminated with eye movements or blinks were not analysed, as these could have affected the deployment of covert attention to target stimuli. Trials with extreme response times (either < Q1 - 2.5*IQR or > Q3 + 2.5*IQR; Q: quartile, IQR: interquartile range) were also excluded from the analysis.

Target stimuli presented on the right and left hemifields were analysed separately to account for potential lateralization effects. We statistically tested the effect of Distractor (positive, neutral, negative, and non-distractor) x Hemifield (right and left) by means of a 4 × 2 ANOVA. In the case of non-sphericity, the multivariate approximation (Wilks’ lambda) was taken into account. Effect sizes were estimated using the partial eta-square (*η*^2^_p_) method. Post-hoc pairwise t-test analyses were conducted to detect specific differences among conditions, adjusting p-values by means of the Bonferroni correction when multiple comparisons were performed. In this case we used the unbiased Cohen’s d (also referred to as Hedge’s *g*) to measure the effect size (Cumming, 2013).

### Recording of the EEG signal

The EEG signal was acquired using an electrode cap (custom-made by ElectroCap International) with tin electrodes. Fifty-nine electrodes were placed at the scalp surface following the 10-10 international system. All electrodes were referenced to the nose-tip and grounded with an electrode located on the forehead. Additionally, horizontal and vertical electro-oculographic (EOG) activity was recorded to monitor blinks and eye movements. Recordings were continuously digitized at a sampling rate of 420 Hz and on-line bandpass filtered between 0.3 and 1.000 Hz.

### Analysis of the EEG signal

EEG data analysis was carried out using the FieldTrip toolbox (version 20180405; Oostenveld et al., 2011) and in-house Matlab code. Overall, we aimed to test whether emotionally salient distractors modulate alpha oscillations, similarly to the modulation of alpha-band power induced by endogenous attention. To this end, we first pre-processed and removed artifacts from the EEG signal. We then performed a time-frequency (TF) analysis to identify frequency bands and electrodes of interest. Once these were established, we computed and statistically compared the temporal-spectral evolution (TSE) curves in response to different distractors and in the absence of distraction. These analysis steps are described in more detail in the following sections.

#### Pre-processing

The EEG signal was segmented into 6-s-long trials (3.5 s pre-stimulus) time-locked to both cue and target/distractor onset. Long epochs were used to avoid edge artifacts in subsequent spectral analyses. Each trial was assigned to one of the four experimental conditions: positive, neutral, negative, or no distractor.

We then carried out a three-step artifact rejection procedure. First, trials contaminated with eye movements were eliminated, as these may indicate a failure to maintain covert attention. Second, any remaining artifacts were removed by means of independent component analysis (ICA) on a 20-dimensional space obtained by principal component analysis (PCA). Finally, noisy channels were interpolated based on data recorded on adjacent electrodes.

One participant was not included in the EEG data analysis due to an excessive number of ocular artifacts (> 70% of trials contaminated). The average number of artifact-free trials per condition in the remaining sample was 80.9 ± 14.3 and did not significantly differ across conditions (F_3,102_ = .514, p = .597).

#### Time–frequency (TF) analysis

TF analysis aimed to identify the frequency range and electrodes with power modulations. Spectral power was computed for each trial using a Hanning-tapered sliding window Fourier transform from 0 to 20 Hz in 0.5-Hz steps and from −0.5 to 1 s in 50-ms steps. The width of the Hanning window was set to 0.8 s to maximize spectral resolution. Power was normalized by computing the relative percentage change from baseline. In order to determine frequency bands and channels of interest, we computed the average oscillatory activity contralateral to both target and distractor presentation for all conditions together.

#### Temporal–spectral evolution (TSE) curves

TF analysis revealed power modulations centred around 11 Hz and maximal over O1/O2 electrodes. Based on this, we computed the TSE of alpha-band oscillations time-locked to cue and target/distractor onset for these electrodes. To do so, we bandpass filtered single-trial EEG activity between 9 and 13 Hz and subsequently calculated the absolute value of the Hilbert transform. TSE curves were baseline corrected using a 0.5 s baseline segment.

Contra and ipsilateral TSE curves time-locked to distractor appearance were statistically compared with trials on which no distractor was present. Each distractor condition (positive, negative, neutral) was tested separately by means of a non-parametric, permutation-based approach. In each permutation, trials of the distractor and non-distractor conditions were randomly shuffled and subjected to a paired t-test. This process was repeated 1000 times. To correct for multiple comparisons, the maximum t-value across time was stored in each repetition. The resultant distribution of the t-statistic was used to derive corrected p-values. T-values above the 95^th^ percentile were considered to be statistically significant.

### Correlation of alpha-band power with behavioural performance

We then tested whether modulation of alpha-band activity was correlated with behavioural performance. For this purpose, we calculated the variation in accuracy in the presence of distractors compared with the non-distractor condition (i.e., distractor-non-distractor). Thus, an improvement in performance under distraction is indexed by positive values, whereas negative values are taken to indicate a deterioration in performance. We then computed the correlation coefficient between accuracy indices and alpha-band power.

### Correlation between alpha-band power and self-reported arousal and valence

Since the largest attentional effects were found for negative distractors, and these tended to be rated as most arousing in the self-report assessment, we conducted an additional analysis to test whether alpha-band suppression might be related to emotional arousal. Correlation analysis was conducted on a single-trial basis. For each participant, we computed the correlation coefficient between alpha-band power and the self-reported arousal of the image presented on each single-trial. Individual correlation values were then subjected to a one-sample t-test compared to zero, to test whether alpha-band modulations and stimulus arousal were either positively or negatively correlated among individuals. Additionally, we carried out the same procedure to assess whether alpha power and emotional valence were correlated at the single-trial level.

## Results

### Self-reported emotional valence and arousal

We first checked whether the participants’ assessment of distractor stimuli actually matched the *a-priori* categorization of images as either positive, negative, or neutral. Self-reported emotional valence was 1.22 ± .336 for positive, −1.10 ± .364 for negative, and .067 ± .195 for neutral images. These values correspond to the emotional categories established *a priori* and significantly discriminate between them (F_2,34_ = 310, p < .001, *η*^2^_p_ = .948; t_35_ > 17.7, p < .001, d_unb_ > 2.89). Furthermore, self-reported arousal was 1.02 ± .557 for positive, 1.20 ± .318 for negative, and -.039 ± .154 for neutral images. As expected, the main effect of arousal was also significant (F_2,34_ = 203, p < .001, *η*^2^_p_ = .923), being higher for both positive and negative stimuli compared with neutral ones (t_35_> 10.7, p < .001, d_unb_> 1.75). Therefore, we can be confident that the distractors employed in this study were appropriately assigned to the corresponding emotional category.

We then conducted a more fine-grained analysis to test whether positive and negative images differed in absolute valence and arousal, and whether valence and arousal was different from zero for neutral images. Since we conducted 4 comparisons, the Bonferroni corrected significance threshold was set to .012 (.05/4). We verified that positive and negative images did not differ in absolute valence (t_35_ = 1.66, p = .105, d_unb_ = .271). Negative images showed a trend to be assessed as more arousing than positive ones (t_35_ = 2.27, p = .030, d_unb_ = .370), although this comparison did not reach the corrected threshold for statistical significance. We also verified that arousal did not differ from zero for neutral images (t_35_ = 1.52, p = .139, d_unb_ = .247), though emotional valence was indeed significantly higher than zero (t_35_ = 2.75, p = .009, d_unb_ = .448).

### Behavioural results

The ANOVA conducted on the percentage of correct responses in the digit-parity task showed neither a significant main effect of Hemifield (F_1,35_= .407, p = .528, *η*^2^_p_= .011) nor a significant interaction between Hemifield and Distractor (F_3,105_ = .117, p = .937, *η*^2^_p_ = .003). In contrast, the main effect of Distractor did result significant (F_3,33_= 3.54, p = .025, *η*^2^_p_= .243). Subsequent comparisons aimed to identify pairwise differences among distractor types. As we performed 6 comparisons, the corrected threshold for significance was .008 (.05/6). Accuracy on the digit-parity task showed a trend to be lower when negative distractors were present compared with both positive (t_35_ = 2.35, p = .025, d_unb_ = .383) and non-distractor (t_35_ = 2.24, p = .032, d_unb_ = .364) conditions. Similarly, percentage of correct responses also tended to be lower under the presence of neutral in comparison with positive images (t_35_ = 2.13, p = .040, d_unb_= .347), although none of these comparisons reached statistical significance when correcting for multiple comparisons (Figure 2).

**Figure 2.**
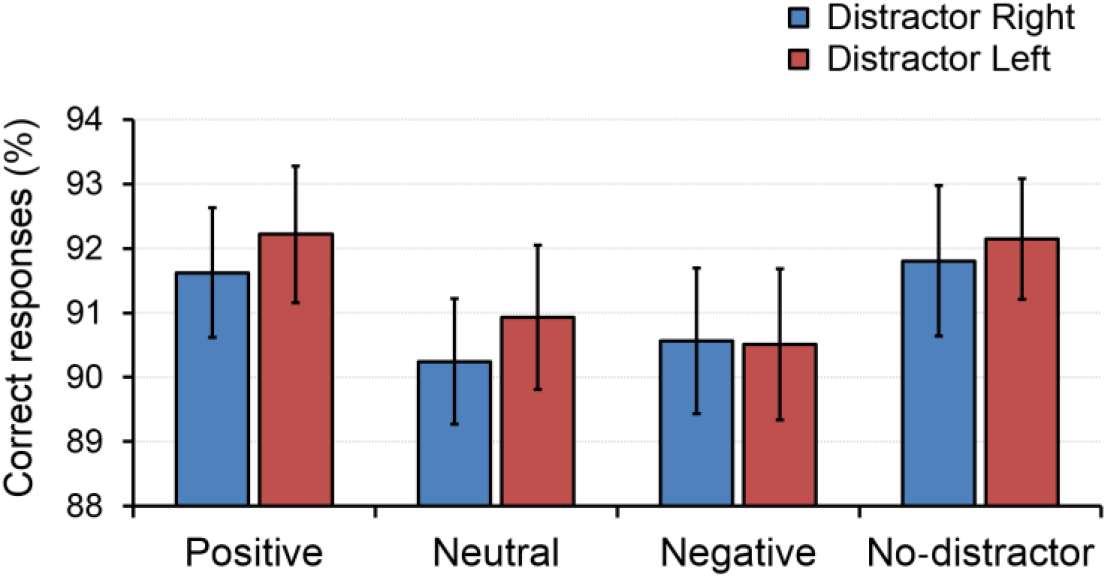
Behavioural results. Percentage of correct responses in the digit-parity task under the presence of positive, neutral, and negative images, as well as in the absence of distraction. Blue bars indicate distractors presented on the right; red bars indicate left. Error bars represent the standard error of the mean.

The ANOVA performed on response times revealed no significant effects (all p > .246) and small effect sizes (*η*^2^_p_ < .116).

### Time–frequency (TF) results

As can be seen in Figure 3, TF analysis revealed a modulation of power in the alpha frequency range in response to both cue and target/distractor presentation. As expected, following cue onset, alpha-band power decreased over contralateral posterior sites, reflecting the deployment of endogenous attention to the cued location. Alpha-band activity was also suppressed in response to the simultaneous presentation of target and distractor, though more bilaterally. Alpha power reduction occurred mainly within the frequency range of 9 to 13 Hz, peaked between 0.3 and 0.7 s after target/distractor onset, and was maximal over occipital electrodes (i.e., O1/O2).

**Figure 3.**
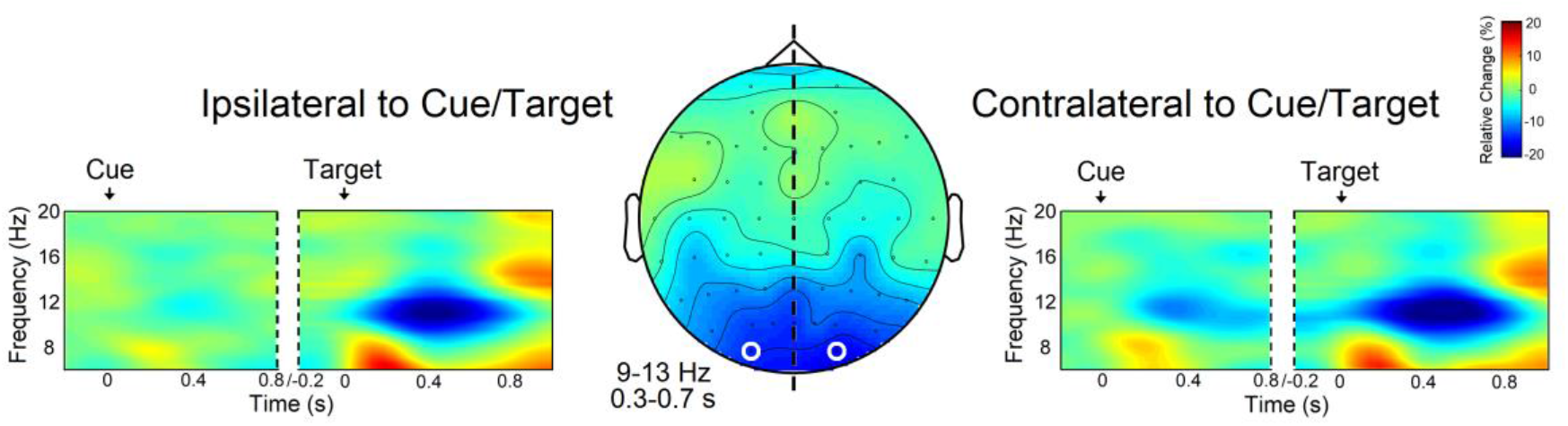
TF results: alpha-band power suppression time-locked to cue and target/distractor. In response to the cue, alpha power decreased over contralateral posterior sites, whereas in response to target/distractor, alpha-band exhibited a bilateral power reduction. The topography shows the modulation of alpha activity between 9 and 13 Hz and from 0.3 to 0.7 s after target/distractor onset. Occipital electrodes with maximal power suppression are highlighted in white.

### Temporal–spectral evolution (TSE) of alpha-band oscillations

Once we had identified the specific frequency band (9-13 Hz) and electrodes (O1/O2) exhibiting maximal alpha-band modulation, we calculated the TSE of alpha oscillations relative to cue and target/distractor presentation. As expected, the allocation of endogenous attention to the cued location was characterized by a reduction in alpha-band amplitude, which was more pronounced contralateral to the focus of attention and peaked at 331 ms (see Figure 4A).

**Figure 4.**
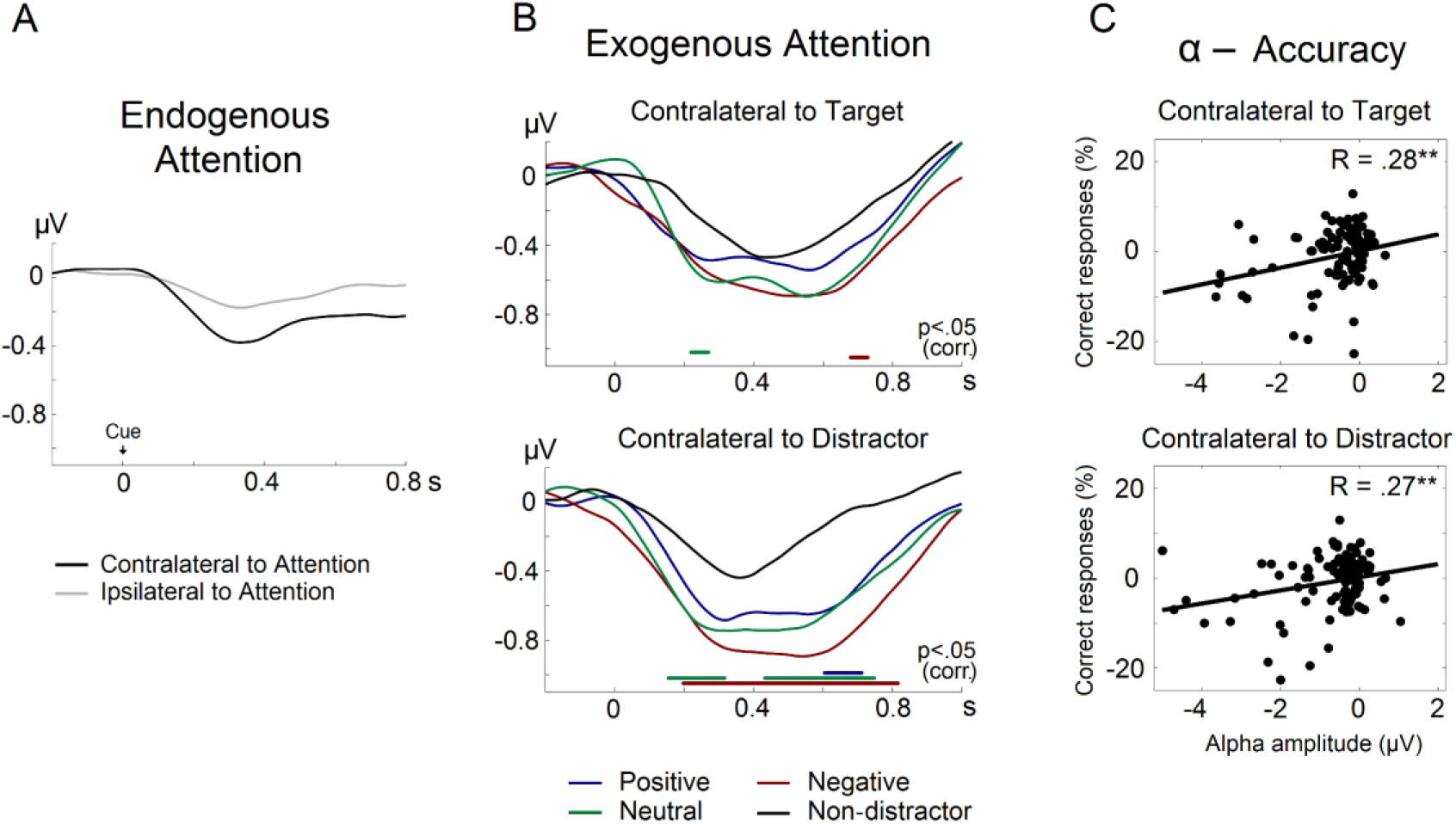
Alpha-band TSE and correlation with accuracy. **A)** TSE of alpha-band amplitude time-locked to cue onset. The black line represents alpha-TSE in the hemisphere contralateral to the focus of endogenous attention; the grey line represents the ipsilateral hemisphere. **B)** TSE of alpha-band amplitude time-locked to target/distractor onset. Alpha-TSE contralateral to target and distractor location are shown separately. The figure shows a more pronounced alpha-band desynchronization contralateral to the presentation of distractor images (positive, neutral, and negative) compared with the absence of distraction. Significant differences between distractor and non-distractor conditions are indicated by coloured horizontal lines (p < .05 corrected for multiple comparisons). **C)** Correlation between alpha-band amplitude and accuracy in the digit-parity task in the hemisphere contralateral to target and distractor. **p < .01

Exogenous attention captured by distractors followed similar temporal dynamics, with a reduction of alpha amplitude reaching a minimum at 329 ms. Statistical analysis revealed significant differences in alpha-TSE depending on whether or not distractors were present. As shown in Figure 4B, distractor effects were mainly evident in the hemisphere contralateral to distractor location. In this hemisphere, alpha amplitude exhibited a higher reduction in the presence of negative images in comparison with the control condition between 198 and 817 ms, for neutral images in two intervals, 155-317 ms and 433-748 ms, and for positive images between 605 and 712 ms. In the hemisphere ipsilateral to distractor presentation two marginal time intervals were found to be significant for negative (679-729 ms) and neutral images (220-270 ms) compared with the absence of distraction (all p < .05 corrected).

### Correlation of alpha-band power with behavioural performance

Correlation analysis was conducted to check for the existence of a relationship between alpha-band power and accuracy in the digit-parity task. In particular, we sought to test if alpha-band modulation was related to the level of attention/distraction devoted to the ongoing task. Analysis was carried out on the time window exhibiting effects in the former TSE analysis (i.e., 0.2 to 0.8 s after target/distractor onset). Our results showed that alpha-band activity is directly correlated with behavioural performance. Thus, the lower the alpha rhythm amplitude, the worse the accuracy (see Figure 4C). Although one would expect this effect to be lateralized, this was not the case, as both alpha-band amplitude in the contralateral (R_103_ = .27, p = .006) as well as in the ipsilateral hemisphere (R_103_ = .28, p = .004) were significantly correlated with accuracy, indicating that the distraction level indexed by alpha-band amplitude results in impaired behavioural performance.

### Correlation between alpha-band power and self-reported arousal and valence

Finally, we conducted a correlation analysis to rule out the possibility that the observed effects could be explained by the arousal of distracting images and not only by their emotional valence. The results of this analysis allowed us to discard this explanation, since the correlation between arousal and alpha-band activity was not statistically significant for either the contralateral (t_34_ = .516, p = .609, d_unb_ = .085) or the ipsilateral (t_34_ = 1.46, p = .153, d_unb_ = .242) hemisphere.

On the contrary, single-trial correlation analysis of self-reported valence and alpha power revealed that lower valence values (i.e., negative distractors) were associated with greater alpha-band desynchronization. This effect was statistically significant for the hemisphere contralateral to distractor location (t_34_ = 2.81, p = .008, d_unb_ = .464), but not for the ipsilateral (t_34_ = 1.72, p = .094, d_unb_ = .284), thus suggesting that the more negative the valence, the greater the capture of attention triggered by emotional distractors.

## Discussion

In the current study, we investigated the neural oscillatory dynamics underlying the attentional capture by emotional stimuli. To this end, we recorded EEG activity in participants performing a visuospatial cueing task, in which a target stimulus appeared in the cued location while an image with either positive, negative, or neutral emotional content was simultaneously presented in the opposite hemifield. Our results show that distractor stimuli, particularly those charged with negative emotion, induce a contralateral reduction in alpha power that is correlated with impaired behavioural accuracy.

As expected, we observed a contralateral decrease of occipital alpha power following cue onset, thus replicating the common finding that lateralized alpha oscillations index the deployment of visuospatial endogenous attention (Capilla et al., 2014; Gould et al., 2011; Thut et al., 2006; van Diepen et al., 2016). Lateralization of pre-stimulus alpha has also been found to be reinforced by increasing the predictive validity of the cue (Dombrowe & Hilgetag, 2014; Gould et al., 2011; van Ede et al., 2012). Hence, by setting the cue validity to 100%, we sought to maximize the expectation of forthcoming target/distractor stimuli and corresponding neural oscillatory effects. In response to the simultaneous presentation of both target and distractor, the pattern of post-stimulus alpha suppression became bilateral over occipital electrodes. Alpha-band activity was reduced contralateral to the location of the target, which is commonly reported when visual stimuli are perceived (Babiloni et al., 2006; Bareither et al., 2014; Harris et al., 2020; Vanni et al., 1997) and has recently been attributed to the attentive processing of target information (Bacigalupo & Luck, 2019; van Diepen et al., 2016). In addition, we found an alpha power suppression contralateral to distractor location, an effect that was particularly marked for images with negative emotional content.

Several authors argue that increases in alpha-band amplitude ipsilateral to the attended visual field are indicative of reduced processing of distractors (Foxe & Snyder, 2011; Jensen & Mazaheri, 2010; Klimesch, 2012; Payne et al., 2013; van Diepen et al., 2016; Zumer et al., 2014). Consequently, higher levels of alpha-band power are thought to reflect a state of inhibited neural processing (Foxe & Snyder, 2011; Jensen & Mazaheri, 2010). However, alpha power enhancement also appears in the absence of distractors (Rihs et al., 2007, 2009; van Gerven & Jensen, 2009) and its role in active distractor inhibition has recently been questioned (Fodor et al., 2020; Foster & Awh, 2019; Schroeder et al., 2018). Indeed, Foster and Awh (2019) speculated whether the alpha power increase contralateral to distractors could simply reflect a by-product of the selection of relevant information rather than suppression of distractors. In an attempt to solve this controversy, it has recently been suggested that target selection and distractor suppression may be two independent processes, both of which are characterized by modulations of alpha oscillations in different regions of the cortex (Wöstmann et al., 2019).

In the present study, the lack of alpha power increase in response to distractor stimuli suggests that these are not being suppressed. Instead, we observed a reduction of post-stimulus alpha-band activity contralateral to distractor presentation. This is clearly reminiscent of the endogenous attention mechanism characterized by an alpha-band desynchronization contralateral to the attended location (Capilla et al., 2014; Rihs et al., 2009; Siegel et al., 2008; Thut et al., 2006; van Diepen et al., 2016) and suggests that attentional resources were allocated to distractor location. In agreement with our results, recent research has also found that exogenous attention driven by non-predictive auditory cues is indexed by an alpha power reduction contralateral to the cued location (Feng et al., 2017; Störmer et al., 2016; but see Weise et al., 2020). Our findings thus support the notion that the involuntary orientation of attention to distractor stimuli is characterized by a desynchronization of alpha oscillations, which would facilitate distractor processing.

Moreover, alpha-band activity has been proposed to index the strength of competition between target and distractor stimuli for attentional resources (van Diepen et al., 2016). Emotional stimuli are biologically salient by definition and are known to capture exogenous attention to a greater extent than neutral distractors (for a review see Carretié, 2014). Therefore, they can be considered as natural competitors against target stimuli for attentional resources. Surprisingly, however, we found no studies employing emotion-laden stimuli to investigate oscillatory correlates of exogenous attention. It has only recently pointed out that the need for proactive and reactive control of emotional distractors is implemented through the modulation of alpha-band oscillations (Murphy et al., 2020), which might also implicitly reflect the degree of attentional capture.

Here, we found that alpha desynchronization was maximal when exogenous attention was drawn to negative distractors. This could be taken to reflect a “negativity bias” in attention, in which negative distractors elicit higher indices of attentional capture than positive and neutral stimuli, as previously reported (Horstmann et al., 2006; Huang et al., 2011; Sussman et al., 2013) and supported by several neural indices (for a review see Carretié et al., 2009). Likewise, event-related potential studies have also revealed that exogenous attention is initially grabbed by images with negative and threatening content (Carretié et al., 2004; Soares et al., 2017). The consequences of ignoring negative stimuli that could signal potential dangers are usually more critical for survival than neutral or even appetitive stimuli, so this is clearly advantageous in evolutionary terms.

Noteworthy, post-stimulus alpha amplitude in response to distractors began to differ from distractor-absent trials as early as 155 ms. This early effect in the modulation of alpha-band oscillations has been to some extent observed in response to exogenously attended stimuli (Feng et al., 2017; Keefe & Störmer, 2021; Störmer et al., 2016). It has been pointed out that exogenous attention to distractors is importantly sustained by the magnocellular visual pathway during the initial stages of visual processing (Carretié et al., 2012, 2017). The magnocellular system is a rapid neural route that processes rudimentary information (Derrington & Lennie, 1984) and has been reported to directly access the prefrontal cortex (Bar et al., 2006), the amygdala (Adolphs, 2004), and the insula (Rodman & Consuelos, 1994), all of them pivotal structures responsible for evaluating stimuli and organizing a response. This could account for the fast alpha-band mechanism underlying exogenous attention observed in the present study. Interestingly, when distractors are biologically salient, the preference for magnocellular processing remains at subsequent stages (Carretié et al., 2017). In fact, it has been proposed that magnocellular involvement in emotional stimulus processing might play a key role in the abovementioned negativity bias, since it would facilitate the detection of urgent visual events (Carretié et al., 2007). Therefore, we could speculate that the enhanced alpha desynchronization for negative distractors found here may be underpinned by the magnocellular visual pathway.

It is important to mention that mere exposure to emotion-laden stimuli has also been related to posterior alpha-band desynchronization (Cui et al., 2013; Knyazev et al., 2008; Mennella et al., 2017; Popov et al., 2012, 2013; Simons et al., 2003). In light of our findings, this effect could be explained by an automatic attentional mechanism triggered by emotionally salient stimuli, which would be reflected in the modulation of alpha oscillations. Nevertheless, it could also be argued that alpha-band suppression does not reflect an attentional bias to emotional information but its inherent processing. Additional research is needed in order to further disentangle perceptual from attentional effects.

Emotional arousal has also been demonstrated to critically modulate alpha oscillations (Cui et al., 2013; de Cesarei & Codispoti, 2011; Schubring & Schupp, 2019; Simons et al., 2003). Indeed, differences in attention-related alpha-band power disappear when emotional stimuli are matched in terms of arousal (Simons et al., 2003). With this in mind, we selected positive and negative images to be equated on arousal values, whilst the neutral stimuli were required to have a score close to zero in this dimension. By analysing the assessment of emotional valence and arousal provided by our participants, we verified that the neutral stimuli were judged to be low arousal images.

Nonetheless, and contrary to our expectations, participants assessed the negative images to be slightly more arousing than the positive ones. Hence, in order to rule out the possibility that the obtained results were rather due to an arousal effect, we analysed the correlation between alpha power and arousal score on a single-trial basis. Critically, we found that alpha-band oscillations were not sensitive to arousal, thus allowing us to discard this alternative explanation. The fact that individual participants rated negative images as more arousing than normative scores has also been observed previously (Weinberg & Hajcak, 2010) and could reflect their individual negativity bias. This explanation is even more plausible given the age range of our sample (19 to 28 years), since younger adults tend to be more vulnerable to distraction by negative stimuli and display a stronger negativity bias (Thomas & Hasher, 2006). Therefore, it should be taken into account that generalization of the present results may be limited due to specific characteristics of our sample.

Contrary to the lack of an arousal effect, we did find a significant correlation between alpha-band amplitude and self-reported ratings of emotional valence at the single-trial level. Thus, the more negative the valence, the lower the alpha-band power. In addition, lower levels of alpha activity were correlated with worse behavioural performance on the ongoing task, which in turn tended to be mostly affected by negative stimuli. Previous research has reported a higher behavioural cost when distractors strongly compete for attentional resources (Fodor et al., 2020; Schroeder et al., 2018; van Diepen et al., 2016). Critically, negative-laden stimuli have been found to impair performance compared with positive (Horstmann et al., 2006; Sussman et al., 2013), as well as with neutral distractors (Horstmann et al., 2006; Huang et al., 2011; Sussman et al., 2013). Taken together, our results support the view that alpha power suppression indexes the degree of attentional capture and subsequent distraction from the ongoing task, which is particularly pronounced in the case of negative distractors.

Interestingly, trials with positive distractors showed a tendency for improved accuracy. It has been shown that positive emotions can broaden the scopes of both attention and cognition, improving goal-directed actions (Fredrickson & Branigan, 2005), an effect that could be mediated by the capacity of positive affect to broaden the field of view in attentional tasks (Schmitz et al., 2009). Further research is needed to elucidate if emotionally positive images may play a protective role in the competition for attentional resources, thus preserving task performance.

In conclusion, we have demonstrated that exogenous attention to emotion-laden visual stimuli is characterized by a pronounced decrease of alpha-band power contralateral to distraction. This effect evolves rapidly and displays a negativity bias, reflecting that negative stimuli may be automatically prioritized in the competition for attentional resources.

## Acknowledgments

This work was supported by FEDER/Ministerio de Ciencia, Innovación y Universidades – Agencia Estatal de Investigación, Spain (grants PGC2018-100682-B-I00 and PGC2018-093570-B-I00) and by the Comunidad de Madrid and the Universidad Autónoma de Madrid, Spain (grants 2017-T2/SOC-5569 and SI1-PJI-2019-00011).

